# Cellular diversity of the somatosensory cortical map plasticity

**DOI:** 10.1101/201293

**Authors:** Koen Kole, Wim Scheenen, Paul Tiesinga, Tansu Celikel

## Abstract

Sensory maps are representations of the sensory epithelia in the brain. Despite the intuitive explanatory power behind sensory maps as being neuronal precursors to sensory perception, and sensory cortical plasticity as a neural correlate of perceptual learning, molecular mechanisms that regulate map plasticity are not well understood. Here we perform a meta-analysis of transcriptional and translational changes during altered whisker use to nominate the major molecular correlates of experience-dependent map plasticity in the barrel cortex. We argue that brain plasticity is a systems level response, involving all cell classes, from neuron and glia to non-neuronal cells including endothelia. Using molecular pathway analysis, we further propose a gene regulatory network that could couple activity dependent changes in neurons to adaptive changes in neurovasculature, and finally we show that transcriptional regulations observed in major brain disorders target genes that are modulated by altered sensory experience. Thus, understanding the molecular mechanisms of experience-dependent plasticity of sensory maps might help to unravel the cellular events that shape brain plasticity in health and disease.

## Introduction

Neurons along the sensory axis in the brain are responsible for the processing and incorporation of inputs originating from the peripheral organs, granting the organism the ability to sense. As the incoming sensory information is often highly complex, the nervous system has to deal with a high-dimensional space in a time-varying manner. For over a century, it has been known that neurons within sensory areas display so-called ‘receptive fields’ which are temporally varying representations of the sensory periphery in individual neurons (Sherrington, 1906). Neurons in the primary visual cortex, for instance, are known to possess orientation selectivity: they respond strongly to visual stimuli of a given specific angle but lose their responsiveness to the same stimulus when it is rotated further away from their preferred angle (Hubel and Wiesel, 1968). Similarly, neurons in the primary auditory cortex respond preferentially to stimuli at a given frequency although their selectiveness diminishes as the loudness of the sound increases (Guo et al., 2012). Sensory cortices are typically organized so that neurons that have similar receptive fields are located in each other’s vicinity thus forming sensory maps in the neocortex (**Figure 1**).

**Figure 1.**
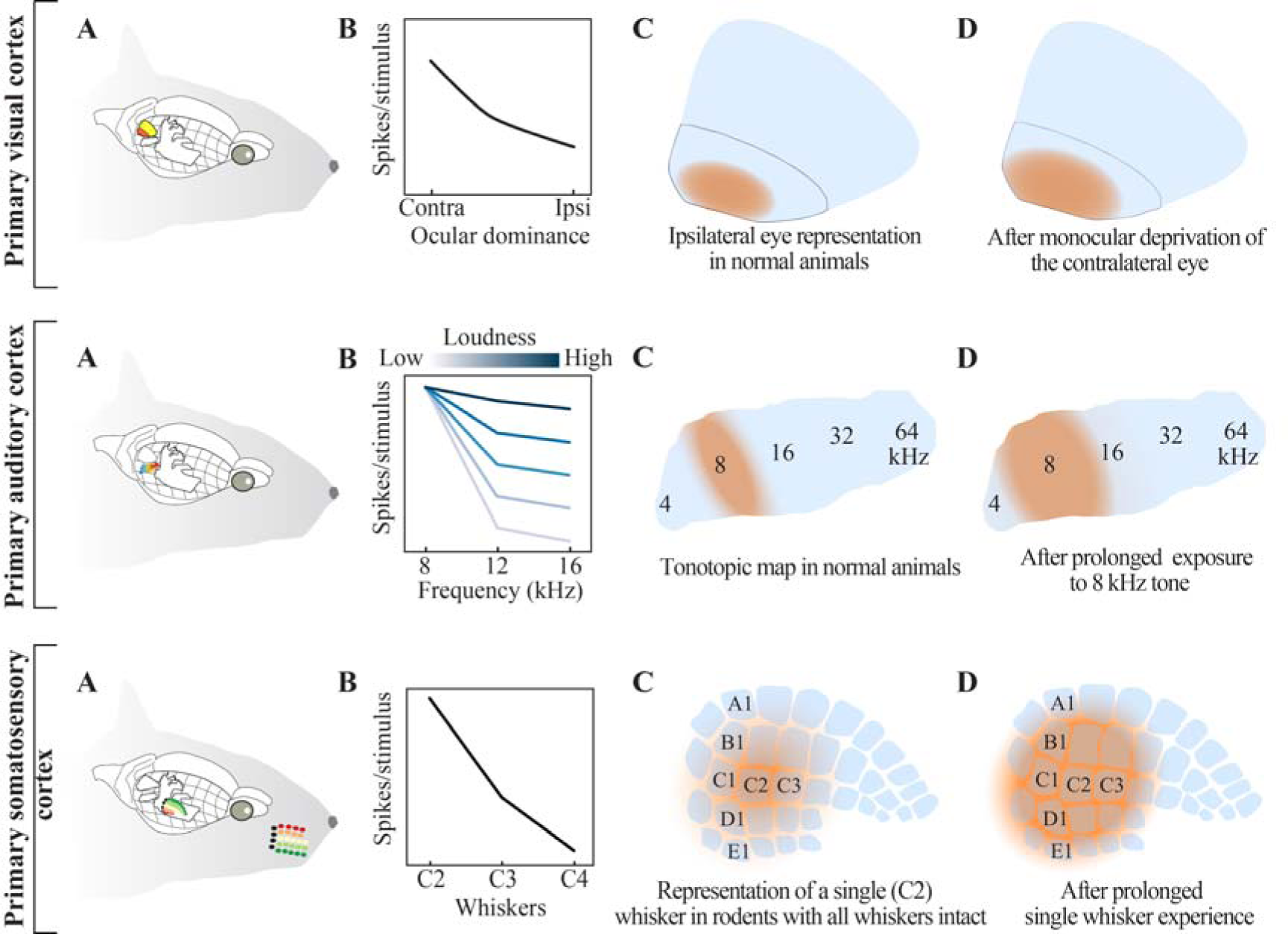
Receptive field plasticity and map reorganization across primary sensory cortices in rodents. (**A**) Relative locations of visual, auditory and somatosensory cortices in the rodent brain; areal designations and locations are approximated. Note that, for clarity, the ipsilateral cortices are shown whereas input mainly originates from the contralateral sensory organs. (**B**) Receptive field organization across the three cortices. Top row: The visual cortex can be subdivided into monocular and binocular region. Middle row: Auditory cortical neurons have a ‘preferred frequency’. As the loudness of the auditory stimulus increases receptive fields are broadened, and neurons respond to stimuli across frequencies. Bottom row: Neurons in the somatosensory cortex are organized in a columnar fashion. Neurons in a column share a common principal whisker to which they preferentially respond. The response amplitude is reduced with increased distance between the principal whisker and the deflected (neighboring) whisker. (**C**) Cortical representations of the ipsilateral eye in the monocular region, a select tonal frequency and a whisker under normal conditions. (**D**) Upon sustained manipulation of sensory input (i.e. monocular deprivation (Gordon and Stryker, 1996), exposure to select tonal frequency(de Villers-Sidani et al., 2007) or whisker deprivation (Fox, 2002)) the sensory maps are reorganized. Independent from the sensory modality, increased use of a sensory organ, or exposure to a specific stimulus, results in expansion of cortical representations in an experience-dependent manner.

Receptive fields are not hard-wired, but adapt to ongoing changes in the statistics of the incoming sensory information. This process, also known as experience-dependent plasticity (EDP), is believed to underlie cortical map plasticity. Experiments across sensory modalities have shown as a general rule that preferential use of a sensory organ, or passive exposure to a select stimulus feature, results in the expansion of the sensory organ, or stimulus representation in the cortex (**Figure 1**). Synaptic and network mechanisms of EDP are increasingly well understood (Feldman, 2009; Feldman and Brecht, 2005; Fox and Wong, 2005; Froemke, 2015), and commonly studied in the context of changes in electrophysiological properties of neurons and synaptically coupled networks. As long-lasting changes in synaptic organization require molecular regulation in individual cells, there is a growing interest in systematic identification of molecular pathways that control EDP to mechanistically understand how experience alters neural networks and shapes behavior. Here, focusing on the map plasticity in the primary somatosensory cortex, we review the state-of-art of molecular correlates of EDP, and perform a meta-analysis of transcriptional and translational changes observed upon altered whisker use. We argue that experience alters gene regulation not only in neurons but in other types of cells in the brain, although the time-course, the dynamics (e.g. up- vs downregulation) and the pathways of plasticity are at least partially cell-type specific. Linking the synaptic activation in neurons and the subsequent regulation of releasable molecules to changes in neurovasculature, we further address how systems level brain plasticity can be modulated.

### 1) The whisker-barrel system as a model system to study neuroplasticity

Cortical representations of rodent whiskers in the barrel cortex, a subfield of the somatosensory cortex (Van der Loos and Woolsey, 1973) have become one of the leading animal models to study experience-dependent plasticity.They possess several distinct advantages over other models of EDP, including:(1) whiskers are organized in an orderly manner on the rodent’s snout, with ~32 macro vibrissae (i.e. whiskers) spanning across 5 rows (named A-to-E; color coded on the figurine) and 4-8 arcs, and topographically represented by neighboring cortical columns in the barrel cortex (**Figure 1C, bottom**). This discrete topographic map allows easy and reproducible identification of the neural circuits that are altered by differential sensory organ use. (2) just like fingers they are represented as somatosensory and motor maps in the cortex, but unlike fingers, whiskers can be reversibly deprived, providing a unique opportunity to study map reorganization during sensory deprivation and recovery from sensory organ loss. (3) whisker deprivation can be applied with ease and is relatively mild compared to other sensory deprivation paradigms such as ocular closure or digit amputation, and to some degree belongs to the rodents’ natural sensory experience. Whiskers, like other hairs, undergo a hair-cycle, which results in spontaneous whisker loss, and rodents often pull their own whiskers while grooming and commonly barber each other’s whiskers during social interaction (Sarna et al., 2000). Whisker deprivation is therefore likely to be less stressful for experimental animals, compared to the other established deprivation protocols across sensory modalities. Whisker plucking or clipping also differs markedly from follicle ablation, ocular removal, cochlea removal, follicle lesioning or nerve severance, which not only irreversibly eliminate peripheral sensory input, but also lead to nerve degeneration, nerve regeneration and cell death, which on their own could influence neuronal responses (Hámori et al., 1986; Waite and Cragg, 1982). These characteristics make the rodent whisker-to-barrel system ideally suited for the study of cortical EDP and as such it is widely used for this purpose.

### 2) Types of receptive field plasticity in the barrel cortex

As originally shown in the barrel cortex (Hand, 1982), sensory deprivation induced by transient whisker trimming is sufficient to perturb receptive field organization both during development and in adulthood (Kossut, 1985; Land and Simons, 1985). The extent of plasticity depends on the nature and duration of sensory deprivation as well as the age of the animal. Synaptic competition for sensory input is hypothesized to be a major driving force for cortical map plasticity (Fox, 2002; Song et al., 2000) and can readily be introduced through a wide range of deprivation protocols. Trimming all whiskers, but one, for example, results in expansion of the spared whisker’s representation (**Figure 2B**) (Fox, 1992) while depriving all whiskers except two neighboring ones, fuses the representation of the two spared whiskers (**Figure 2C**) (Diamond et al., 1994). Other methods of deprivation range from single or multiple row sparing(Broser et al., 2008; Finnerty et al., 1999; Kaliszewska et al., 2012) or deprivation(Allen et al., 2003) and single whisker deprivation (**Figure 2D**) (Celikel et al., 2004) to more complex deprivation protocols (e.g. chessboard pattern deprivation, where every other whisker is deprived (Trachtenberg et al., 2002; Wallace and Fox, 1999)). Independent from the type of the deprivation protocol employed, these studies commonly concluded that experience-dependent changes in receptive field plasticity can be summarized by increased representation of the spared, and decreased representation of the deprived whisker. However, receptive field plasticity can also be induced by allowing animals to explore novel (or otherwise enriched) environments (Polley et al., 2004) (**Figure 2E**) or simply by passive whisker stimulation (Welker et al., 1992) (**Figure 2F**). These observations suggest that there are multiple forms of receptive field plasticity that could be modulated by contextual and top-down processes even in the absence of altered sensory organ use.

**Figure 2.**
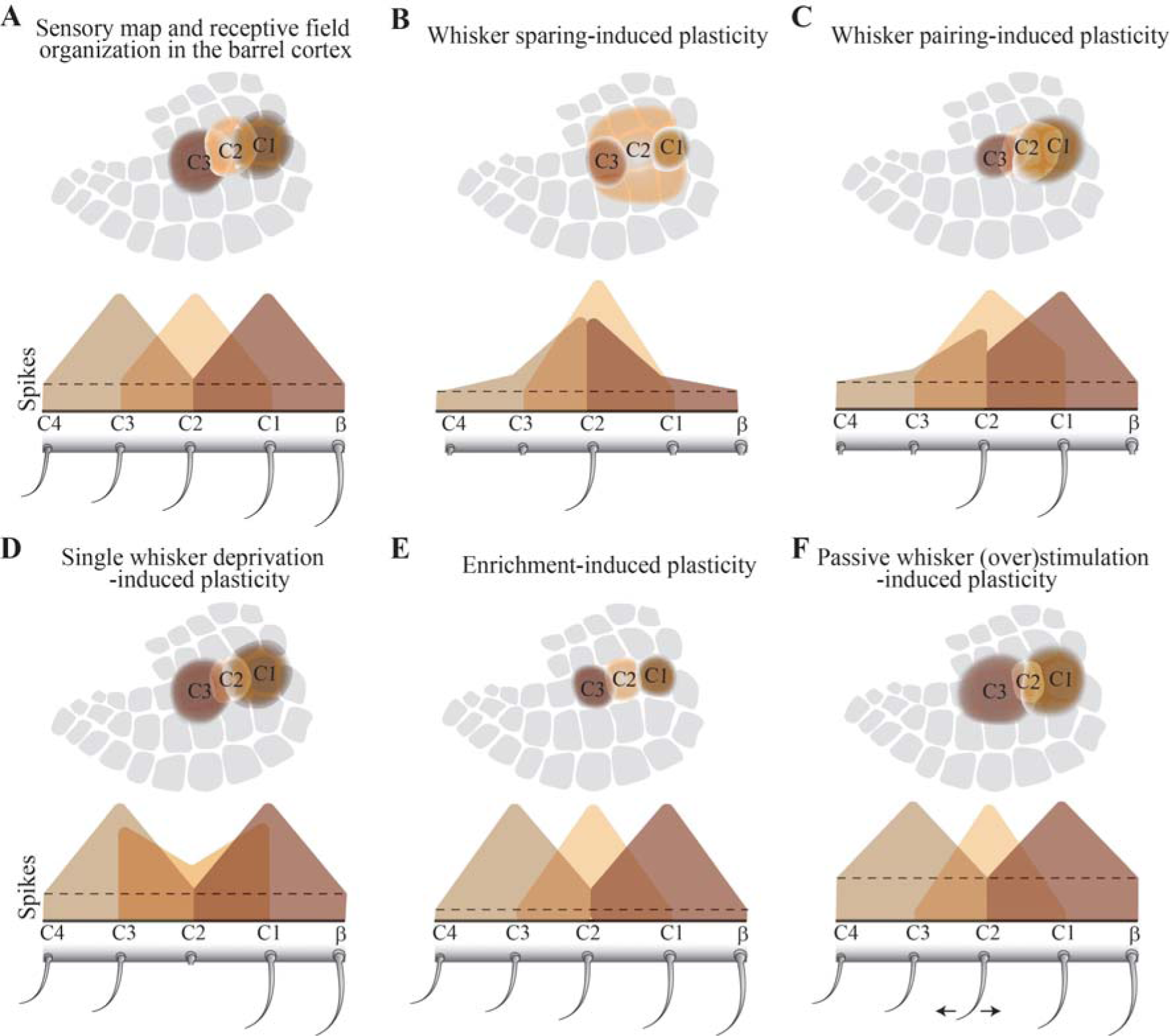
Known types of plasticity in the barrel cortex. Schematic representation of the sensory deprivation/experience condition, representative single neuron receptive fields in the supragranular layers of the barrel cortex, and the presumed map organization in the same layers across five different plasticity induction protocol. (**A**) Neurons in the barrel cortex respond to deflection of multiple whiskers. The whisker that evokes the largest number of action potentials is named as principal whisker, and the others are referred to as surround whiskers. In the absence of any whisker deprivation, perceptual training or environmental enrichment, the receptive field organization follows a topographical mapping where neurons in neighbouring cortical columns have neighbouring whiskers on the periphery as their principal whiskers. Neighboring whisker representations only partially overlap preserving the topographic mapping of the sensory periphery at the level of cortical circuits. (**B**) When all whiskers, but one, are removed, neural responses to the spared whisker are potentiated in neighbouring barrel columns(Fox, 1992), resulting in spared whisker representation to expand into the neighbouring cortical columns whose principal whiskers have been deprived. (**C**) Sparing two neighbouring whiskers causes their receptive fields to merge (Diamond et al., 1994), as spared whisker representations’ increasingly overlap. (**D**) Depriving one (row of) whiskers result in neurons in the deprived cortical column to acquire a new principal whisker (Celikel et al., 2004); while deprived whisker representation shrinks, the spared surrounding whisker representations expand to drive stimulus evoked representations in the deprived whisker’s column. (**E**) Enriched environment experience leads to sharpened receptive fields and whisker representations in the barrel cortex (Polley et al., 2004). (**F**) Chronic (over)stimulation of a single whisker results in a shrunken receptive field in neurons of the corresponding barrel column while the receptive fields of the non-stimulated whisker tend to broaden (Welker et al., 1992). Single whisker representations are believed to expand upon non-stimulated whiskers, and shrink after stimulated whisker’s deflection, reflecting increased topographic precision of the sensory map.

The majority of the studies on receptive field plasticity (and in brain plasticity in general) have focused on neurons. Glial cells can be found in high numbers in the mammalian brain and are indispensable for proper brain function (Azevedo et al., 2009; Jäkel and Dimou, 2017). Neuroglial cells include astrocytes, microglia and oligodendrocytes, which together are involved in, for example, myelination, neurotransmitter recycling, response to brain damage and pathogens, and neurovascular coupling (also see the section on “*From synaptic activity to vascular plasticity”*). Glial cell morphology and activity can be modulated in an experience dependent manner (Barrera et al., 2013; Bergles and Richardson, 2015; Mangin et al., 2012; Stogsdill and Eroglu, 2017). For example, astrocytes’ morphology and abundance are altered upon environmental enrichment, while the number of astrocytic contacts with synapses increase (Black et al., 1987; Diamond et al., 1964; Jones and Greenough, 1996). As astro-neuronal cannabinoid signaling is critical for long-term depression of cortical synapses (Min and Nevian, 2012), sensory deprivation induced synaptic depression might involve both neuronal and glial processes. Both oligodendrocyte morphology and axonal myelination by oligodendrocytes are modulated by recent sensory experience; social isolation or disruption of oligodendrocyte neuregulin signaling reduce axonal myelination and cognitive performance in select long-term memory paradigms (Makinodan et al., 2012); conversely, activity-dependent myelination and oligodendrogenesis improve task performance (Gibson et al., 2014). Microglia are classically known to be involved in the defense against pathogens, but are also involved in synaptogenesis during learning and memory, and respond to sensory input by increasing their contact with dendritic spines and synapses (Parkhurst et al., 2013; Sipe et al., 2016; Tremblay et al., 2010). Given their tight coupling to their neuronal counterparts, it is not surprising that glial cells have a preferred stimulus in the visual cortex (Schummers et al., 2008). Thus, (plasticity of) receptive fields are unlikely to be restricted to neurons, although non-neuronal plasticity in the barrel cortex is not commonly studied.

### 3) Molecular correlates of plasticity

Studies describing the cellular mechanisms underlying experience-dependent plasticity in the barrel cortex system, typically by electrophysiological means, are plentiful (see e.g (Allen et al., 2003; Celikel et al., 2004; Cheetham et al., 2007; Chittajallu and Isaac, 2010; Finnerty et al., 1999; Fox, 1992; Greenhill et al., 2015; Hardingham et al., 2008; Harris and Woolsey, 1981; House et al., 2011; Kätzel and Miesenböck, 2014; Kossut, 1985; Lendvai et al., 2000; L. Li et al., 2009; P. Li et al., 2009; Margolis et al., 2012; Micheva and Beaulieu, 1995; Miquelajauregui et al., 2015; Nicolelis et al., 1991; Simons and Land, 1994, 1987; Stern et al., 2001; Wang and Zhang, 2008; Welker et al., 1989; Zembrzycki et al., 2013)). There has been, however, a surprising lack of experimental studies to unveil the molecular mechanisms that underlie the observed changes in neuronal responses in response to (altered) sensory experience. Single molecules, particularly those that had been previously shown to modulate synaptic plasticity, such as the Ca^2+^/calmodulin-dependent protein kinase (CaMK), the transcription factor Cre-Response Element Binding (CREB), the growth factor Brain-derived Neurotrophic Factor (BDNF) and nitric oxide (Dachtler et al., 2012; Glazewski et al., 1999, 1996; Rocamora et al., 1996) have been studied in some detail, however technologies now allow for systematically addressing the molecular correlates of map plasticity throughout the transcriptome and proteome. Changes in transcripts’ abundance can be quantified through the use of microarrays and RNA sequencing, while the proteome can be surveyed through, for example, tandem mass spectrometry. These techniques provide valuable information on the molecular processes and pathways that are required to establish, maintain and adapt the neural circuit organization in response to sensory experience. Moreover, these data can now also be obtained in cellular and subcellular resolution as individual cells can be sorted and their RNA subsequently sequenced (Zeisel et al., 2015).

#### 3.1) Input-dependent and cell type specific effects of brief enhanced sensory experience on barrel cortex transcriptome

In a pioneering study that employed microarraystoquantitativelyaddress transcriptional regulation in the barrel cortex, Vallès and colleagues studied animals upon exposure to an enriched environment (EEE) (Vallès et al., 2011). Adult rats in the experimental group were subjected to 30 minutes of EEE in a dark room filled with novel objects and toys. The control group was kept in the dark for the same duration but in their own home cages. Animals were sacrificed, either immediately after EEE or following a 4 hour period spent in their home cages, before isolation of barrel cortical RNA which was subjected to microarray analysis, thereby identifying EEE-induced changes in the transcriptome.

A common way to interpret large-scale molecular datasets is through the use of gene ontology (GO) terms. Gene ontologies entail defined cellular components, molecular functions or biological processes to which genes (or rather, their protein products) contribute(Gene Ontology Consortium, 2008). Gene ontology analyses determine whether gene sets that are differentially regulated belong to distinct GO terms and if their co-regulation is significant. GO classification of the differentially expressed transcripts in the Valles et al dataset has shown that gene transcription rapidly changes upon EEE (**Figure 3**), with a time course similar to whisker deprivation-induced regulation of gene transcription (Bisler et al., 2002). The majority of transcriptional regulation after EEE can be linked to general cellular processes (**Figure 3A, Supplemental Tables 1 and 2**), although the direction of the transcriptional regulation, i.e. up- vs down-regulation, depends on the sensory history; immediately after EEE the vast majority of differentially expressed transcripts are up-regulated (170 upregulated, 31 downregulated), whereas in the 4h group downregulated genes are more prevalent (29 upregulated, 98 downregulated). Lack of the sustained upregulation of gene transcription in this group might reflect the fact that animals were returned to their home cage, i.e. an environmentally impoverished environment, before tissue collection for a period of 4h. Expected reduction in the utilization of the somatosensory input to explore animals’ familiar environment might in turn diminish synaptic transmission and, consequently, the rate of metabolism, thus obviating upregulation of transcripts. If an enriched environment modulates gene expression in an activity-dependent manner, one might predict that longer EEE would result in sustained transcript up- or downregulation lasting until the network has accommodated to the enhanced sensory input. This was confirmed in a follow-up study in which rats were subjected to EEE for 28 days, after which only 29 genes were found to be differentially expressed, likely reflecting the ‘steady-state’ of the cortical reorganization upon chronic alteration in incoming sensory information (Vallès et al., 2014).

**Figure 3.**
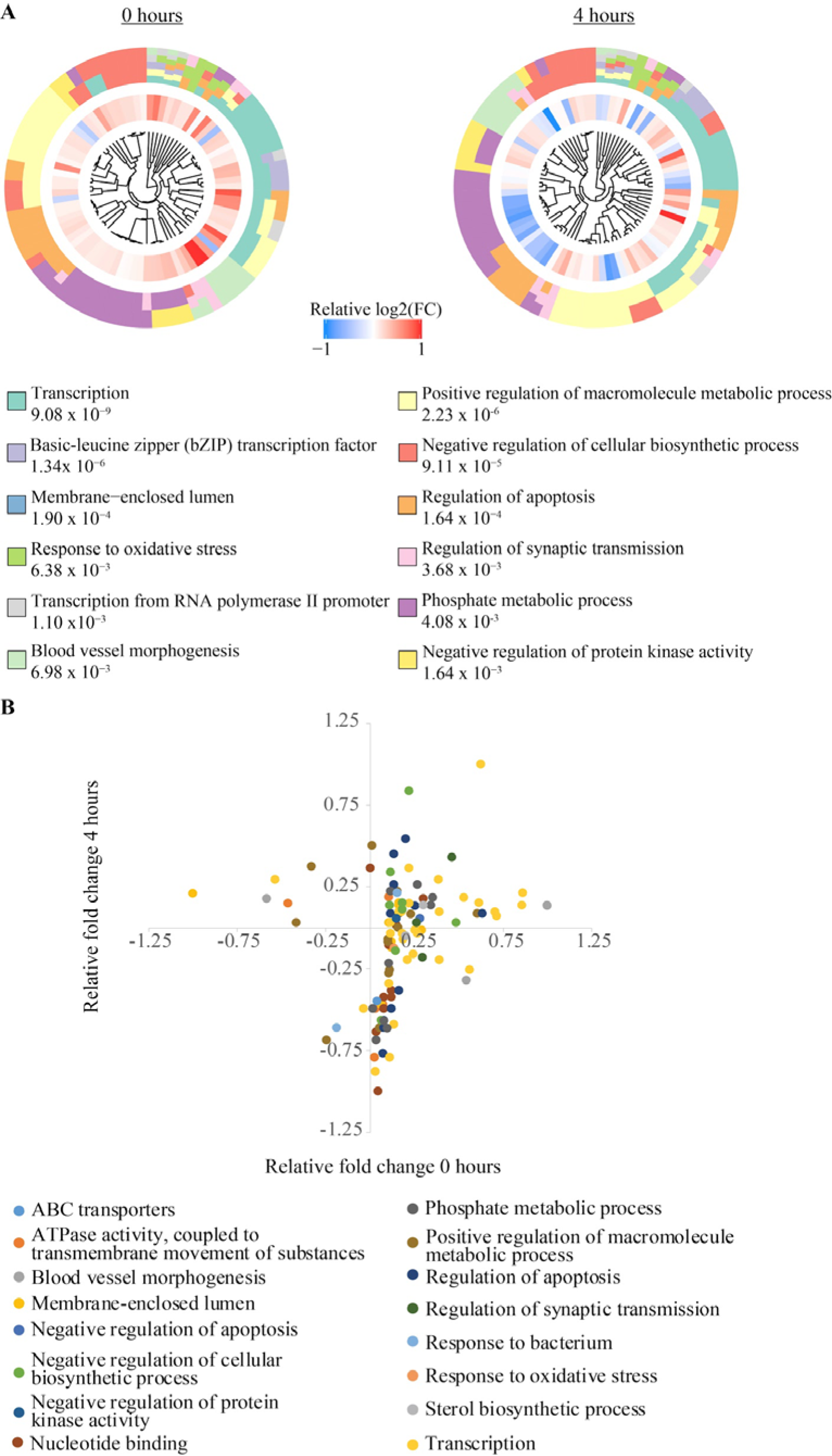
Gene ontology of differentially expressed genes after exposure to an enriched environment. (**A**) Twelve most significant GO terms based on the transcriptional changes after EEE in the barrel cortex (data from (Vallès et al., 2011)). Transcripts are clustered by their respective GO term, numbers indicate p-values. Note that most genes are commonly classified under multiple GO terms. (**B**) Relative (with respect to control condition) expression values of all differentially regulated transcripts at 0 or 4 hours. The color code denotes GO terms. Those transcripts that appear in multiple GO terms are plotted only once; the cluster membership is ranked. As such only the most differentially regulated GO term is displayed for those transcripts that are classified under multiple GO terms. The majority of transcripts (n=103) are upregulated following EEE, half of which remain upregulated in the 4h group (upper right quadrant, n=52), the other half is downregulated (lower right quadrant, n=51). Only few genes have steady downregulation (lower left quadrant, n=2), or temporally delayed upregulation (upper left quadrant, n=6). Thus exposure to an enriched environment triggers temporally varying transcriptional regulation. The direction of change in single gene transcription can be used to classify transcriptional dynamics and relate it to behavioral context.

The overrepresented GO terms identified after EEE are not specific to neurons or synapses. They mostly represent general cellular processes such as transcription, metabolism and cell signaling, which suggests that after EEE, gene expression changes might not be restricted to neurons. Neuronal plasticity might possibly cause, or otherwise contribute to these observed changes in other cell types in the brain. Although the term ‘regulation of synaptic transmission’ (which specifically refers to communication through chemical synapses) was found to be overrepresented, other differentially regulated genes were strongly related to blood vessel morphogenesis. Sustained increased neuronal activity leads to heightened energy consumption and thus elevates oxygen consumption (Shetty et al., 2012). Increased metabolic rate and subsequent hypoxia could also elevate reactive oxygen species, which have been shown to induce oxidative stress and ultimately neuronal death (Friberg et al., 2002). Thus, overrepresentation of GO terms such as ‘response to oxidative stress’ and ‘blood vessel morphogenesis’ point to an important role for vascular endothelial cells in experience-dependent plasticity, serving to accommodate increased energy consumption (also see the section “From synaptic activity to vascular plasticity” below). This also suggests that the transcriptomes of distinct cell types might be differentially regulated by EEE. To explore the possibility, using publicly available single-cell transcriptomics data from experimentally naïve juvenile mouse somatosensory cortex (Zeisel et al., 2015), we calculated a cell enrichment index (CEI; **Figure 4**, **Supplemental Table 3**). The CEI was calculated by dividing the average copy number of each transcript within each cell class by the total copy number of the same transcript averaged across all cell classes. From the list of differentially expressed transcripts identified by Vallès and colleagues (2011), we then selected transcripts that were enriched by >10% (i.e. had a ~10% increased CEI based on the Zeisel dataset) in one cell type compared to the five others, which showed that most differentially expressed transcripts were preferentially expressed in interneurons, whereas astrocytes were the least represented cell class in the Valles dataset (**Figure 5A**).

**Figure 4.**
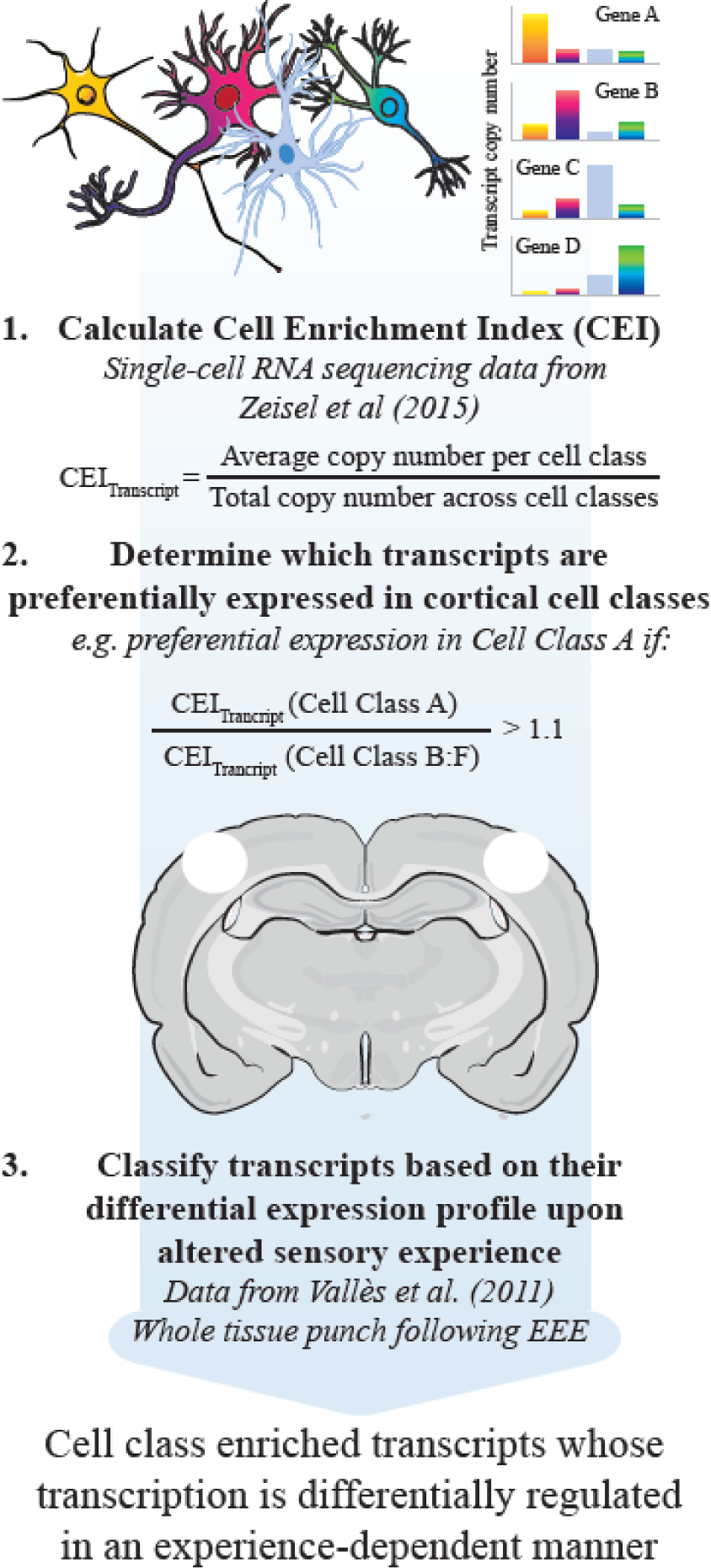
The bioinformatics workflow.

See Supplemental Table 3 for calculated CEIs per transcript.

**Figure 5.**
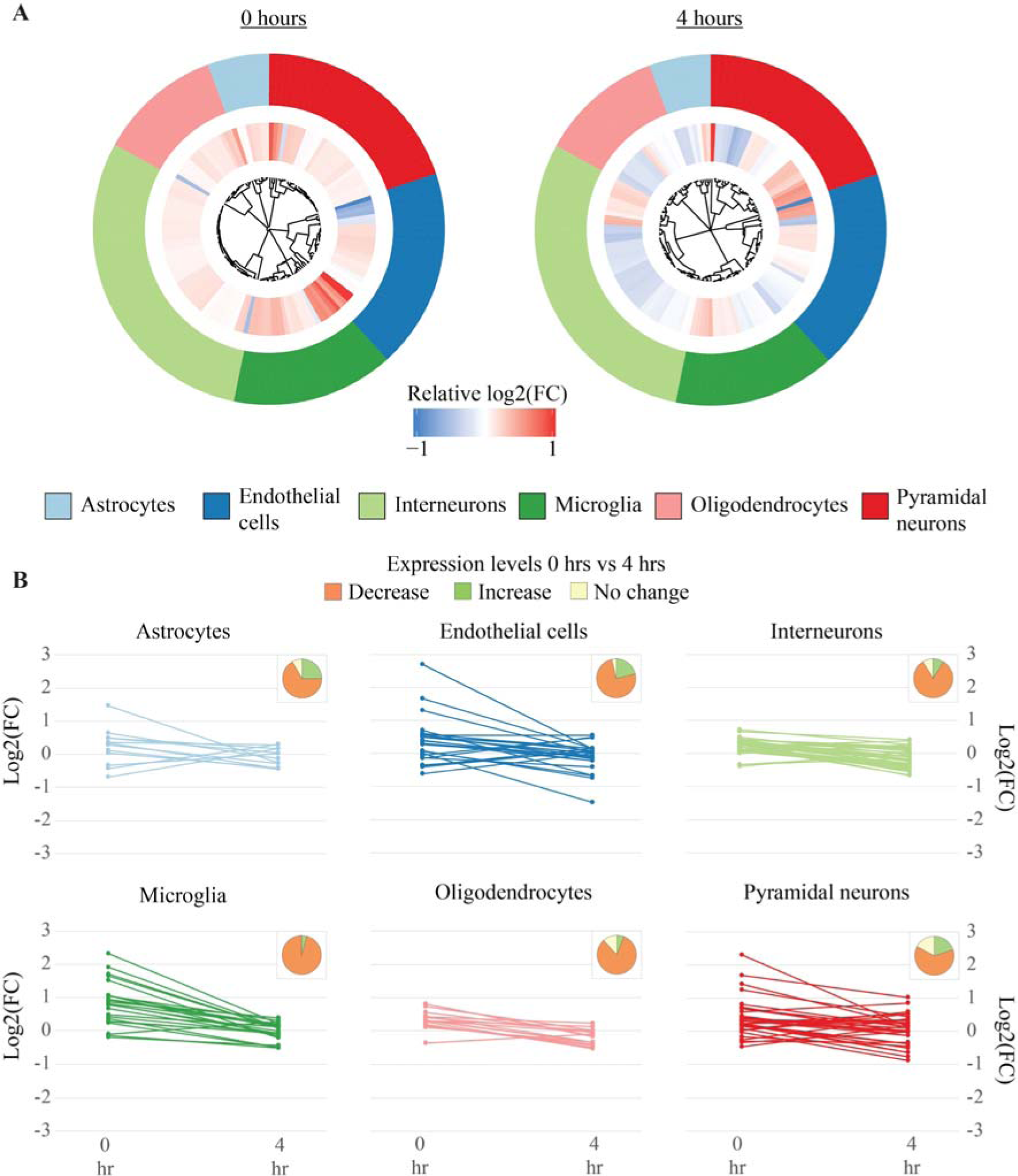
Cell type-specific gene expression profiles after exposure to an enriched environment. Genes that are preferentially expressed in a cell-type specific manner was determined using single-cell RNAseq data ((Zeisel et al., 2015); see main text for details). (A) Distribution of differentially expressed genes 0 or 4 hours after EEE across identified cell classes. (**B**) Temporal changes in gene transcription across cell types. Each data point represents a unique transcript identified as differentially expressed following enriched environment experience (EEE) (Vallès et al., 2011). Pie charts represent the relative percentage of genes whose transcription is downregulated (in orange), upregulated (in green) or non-regulated (in yellow) when the normalized transcription levels in 4h group were compared to 0h.

Genes that displayed the most robust expression changes in response to EEE were preferentially expressed not only in pyramidal cells but also in endothelial cells and microglia. Transcripts found mostly in astrocytes, interneurons and oligodendrocytes, by contrast, showed modest changes after EEE (**Figure 5A,B**). Comparing gene expression across 0h or 4h showed that the majority of transcripts were reduced in their abundance at the 4h interval across all cell types, often returning (close) to baseline levels (**Figure 5B**). These changes were most diverse (albeit only marginally) in pyramidal cells (7 up-, 22 downregulated, 6 no change) while endothelial cells had relative abundance of genes whose transcription is upregulated after 4 hours (**Figure 5B**). Despite the fact that we calculated the CEI based on cells obtained from experimentally naïve juvenile mice and use it to address the plausible cellular diversity in adult rodents, these results argue that sensory experience might affect gene expression in a cell type-specific manner. If so, cellular signaling pathways upstream of differential gene expression are also likely to be distinct for each cell class. A systematic analysis of the mouse transcriptome upon altered sensory organ use will help to unravel the cellular diversity and molecular pathways associated with experience-dependent plasticity.

#### 3.2) Extended sensory stimulation and deprivation and their effects on synaptic proteins

Transcriptional modulation by sensory experience will alter structural and functional organization of neural circuits only if the changes are reflected in the proteome. Because RNA translation into protein is not a linear process, and controlled by posttranscriptional regulatory mechanisms (Keene, 2007), proteomic studies will be required to mechanistically address the molecular pathways associated with expression of plasticity. Currently the only proteomics study available in barrel cortex was performed by Butko and colleagues (Butko et al., 2013), who either clipped or brushed the whisker pads of rats for a period of 30 days, starting from P4, and subsequently prepared synaptoneurosomes from barrel cortex. Using tandem mass spectrometry Butko and colleagues identified systematic downregulation of translation for various ion channels (e.g. inward rectifying potassium channels KCNJ3 and KCNJ6, calcium-activated potassium channel KCNN2), neurotransmitter receptors (e.g. AMPA and NMDA receptor subunits GluA1, GluA2, GLuN1, GluN2A/B), cytoskeletal proteins (e.g. Catenins) and signaling proteins (e.g. adenylyl cyclase, protein kinase C) in whisker deprived rats. These alterations in protein levels are likely to be the cause of electrophysiological changes in receptive field and sensory map representations.

### 4) From synaptic activity to vascular plasticity

If brain plasticity is a systems level response to change, as argued above, there must be molecular players that could couple changes in one cell type to another. For example, changes in neuronal activity might be coupled to neurovascularization, helping to control both the neural activity as well as the dynamics of the blood flow in an experience dependent manner (Kole, 2015).

Short-term changes in local blood flow are associated with neuronal activity and modulated by astrocyte-mediated vasodilation, vasoconstriction (McCaslin et al., 2011; Takano et al., 2006) and cholinergic signaling (Lecrux et al., 2017); inhibitory neurons are also implicated in neurovascular coupling as specific subsets of interneurons can have different effects on vascular tone (Cauli et al., 2004). They can induce vasoconstriction through vasoactive proteins such as neuropeptide Y and somatostatin, whereas vasodilation can be achieved through secretion of vasoactive intestinal polypeptide or nitric oxide (NO), a potent vasodilatory molecule. Dynamic modulation of the cerebrovasculature thus involves a wide range of neuronal and non-neuronal cell types.

Besides short-term adaptation, long-term effects on the neurovasculature also have been observed. Already over two decades ago, in the visual cortex of young and adult rats synaptogenesis was found to be accompanied by angiogenesis (i.e. the formation of new blood vessels) following EEE (Black et al., 1989, 1987). More recent research has shown that during a critical postnatal developmental period, excessive stimulation induces a reduction of blood vessel sprouting and endothelial proliferation in the somatosensory, auditory and motor cortices (Whiteus et al., 2014). These anti-angiogenic effects were associated with hypoxic conditions in the corresponding overstimulated brain regions, which in turn was accompanied dendritic spine loss. A separate study (Lacoste et al., 2014) employed both enhanced and decreased whisker experience, and found that blood vessel patterning in barrel cortex was modulated as a result: vascular density and branching were diminished in response to whisker deprivation whereas the same metrics were enhanced upon whisker stimulation. These findings indicate that the organization of the neurovascular bed is modulated in the wake of sensory experience, and suggest that brain oxygen levels (which are a derivative of brain vascularization) can in fact form a limiting factor to synaptic plasticity.

Despite the clear demonstrations of the impact of sensory experience on the neurovasculature, it is currently unknown which molecular components might orchestrate experience-dependent blood vessel patterning in the neocortex. To establish potential cellular signaling pathways involved in this process, we identified the top ten most differentially regulated genes after EEE (Vallès et al., 2011). We specifically focused on the genes that are in the GO term ‘Vasculature development’ and whose transcript was differentially regulated ≥ 10% beyond control both in 0h and 4h groups (see **Supplemental Table 2 and 4**). These transcripts include *Cyr61*, *Plat*, *Jun*, *Junb*, *Verge*, *Egr1*, *Egr3*, *Nr4a1*, *Nr4a3* and *Hspb1*. The products of these ten genes, discussed in detail below, are involved in a variety of functions that would enable neurovascular plasticity ranging from induction of vessel sprouting and cell migration to alterations in vessel permeability.

#### 4.1) Molecules of neurovascular plasticity

Cysteine-rich angiogenic inducer61 (Cyr61) codes for a secreted extracellular matrix(ECM) protein involved in processes involvingcellmigration, adhesion, differentiation and mitogenesis (Lau and Lam, 1999). In cortical neurons, mechanical strain and BDNF have been shown to induce transcriptional activity of Serum Response Factor (SRF), which in turn is involved in regulating *Cyr61* promoter activity (Hanna et al., 2009; Kalita et al., 2006); A third way that *Cyr61* expression could be regulated by neuronal activity is through muscarinic acetylcholine receptors (mAChRs) and NMDA receptors (Albrecht et al., 2000; Ito et al., 2007).

The tissue plasminogen activator (Plat) is involved in ECM degradation, a critical step in angiogenesis that serves to clear the way for migration of vasculature-associated cells. Its gene product has been shown to be secreted by vascular endothelial cells in response to fluid shear stress, linking its function to neuronal activity resulting in increased blood flow (Diamond et al., 1990). Mechanical stretch induced expression of vascular endothelial growth factor (VEGF) and angiotensin (AGT) requires activation by PLAT (Teng et al., 2012). Its gene product is also involved in mediating synaptic growth, increasing NMDAR-mediated Ca2+ influx and cleaving proBDNF into BDNF and hence also has functions related to neuronal plasticity (Baranes et al., 1998; Nicole et al., 2001; Pang et al., 2004).

Activator protein 1 (AP-1) is a transcription factor regulating various cellular processes such as cell proliferation, differentiation, apoptosis, and cell migration (Angel and Karin, 1991; Karin et al., 1997). It consists of homo or heterodimers of Jun proteins, among which are JUN and JUNB, the transcripts of which were differentially expressed following EEE (Vallès et al., 2011). Inhibition of AP-1 results in reduced vascular smooth muscle cell proliferation and diminished migration and tubule formation by ECs. In addition, inhibited microvascular endothelial cell proliferation and was observed, as well as reduced vasculature formation by ECs after cerebral ischemia (Ennis et al., 2005; Murata et al., 2012; Zhan et al., 2002; Zhang et al., 2004). *Jun* and *Junb* have both been shown to be induced by increased neuronal activity and plasticity (Abraham et al., 1991; Nedivi et al., 1993; Sonnenberg et al., 1989).

The product of the vascular early response gene (Verge or Apold1) accumulates at the periphery of endothelial cells, where it functions to regulate cell permeability by remodeling the cells’ cytoskeleton, allowing substances to pass through endothelial cell layers (Regard et al., 2004). A knockout of this gene is shown to exacerbate outcomes after stroke in mice (Mirza et al., 2013). Conditions inducing its expression has been studied in various tissues, and include hypoxia, ischemia, physical activity and hypertonicity (Liu et al., 2012; Maallem et al., 2008; Regard et al., 2004; Simonsen et al., 2010)). A putative mechanism linking neuronal activity to Verge expression entails tumor necrosis factor (TNF) and fibroblast growth factor (FGF) signaling, which are expressed in response to glutamate and oxygen-glucose deprivation by neurons and astrocytes (Hurtado et al., 2002; Pechán et al., 1993).

Intracellular signaling pathways activate cellular processes downstream of membrane receptors. Prkx codes for a cAMP-dependent protein kinase involved in transcriptional regulation through interaction with and/or phosphorylation of its downstream targets (Zimmermann et al., 1999). Several downstream targets of PRKX are involved in vascular functions such as maintenance of blood vessel integrity and (inhibition of) angiogenesis (Kim et al., 2000; Li et al., 2011; Zhou et al., 2007). Indeed, Prkx expression has been shown to be involved in vascular endothelial cell proliferation, migration, vascular-like structure formation and morphogenesis (Li et al., 2011, 2002). The neuronal function of PRKX is has not been studied in detail, hence as of yet it is unknown whether its regulation can be modulated through increased neuronal activity.

*Egr1* and *Egr3* code for zinc finger transcription factors. They are immediate early genes, as such their expression can be swiftly regulated in response to stimuli. Befitting this characteristic, these two genes are upregulated immediately after EEE (Vallès et al., 2011). They are important for growth factor expression and -signaling; upon *Egr1* knock-down, not only EC replication, migration and tubule network formation was reduced but also expression of FGF was impaired (Fahmy et al., 2003). EGR3 on the other hand is involved in VEGF signaling, as its perturbation is paired with impaired VEGF-induced migration, proliferation and tube formation by endothelial cells (Liu et al., 2008; Suehiro et al., 2010). EGR proteins are upregulated in response to various growth factors (Biesiada et al., 1996; Fang et al., 2013; Mayer et al., 2009; Tsai et al., 2000).

Members of the Nuclear Receptor Subfamily, *Nr4a1* and *Nr4a3* are involved in vascular function through regulation of VEGF signaling (Rius et al., 2006; Zeng et al., 2006), thus helping to control microvessel permeability (Zhao et al., 2011) and endothelial and vascular smooth muscle cell proliferation (Nomiyama et al., 2006; Rius et al., 2006). Nr4a1 also has neuronal functions, as it controls surface expression of the NMDAR subunit NR2B (Zhang et al., 2016) while also regulating spine density in hippocampal excitatory neurons in an activity-dependent manner (Chen et al., 2014). Interestingly, an important upstream regulator of Nr4a receptors, cAMP-response element-binding protein (CREB) (Volakakis et al., 2010), is a key molecule in plasticity induction (Benito and Barco, 2010) but also regulates cellular responses to hypoxia (Leonard et al., 2008; Shneor et al., 2017), suggesting a dual role for the NR4A protein family in neurons as well as the vasculature. Activity-dependent secretion of growth factors, e.g. upregulation of *Nr4a1* and *Nr4a3* upon EEE, could have important functions in, for instance, neuron-to-vasculature signaling (see **Figure 6** and the section *Linking synaptic activity to vasculature change through releasable molecules*)

**Figure 6.**
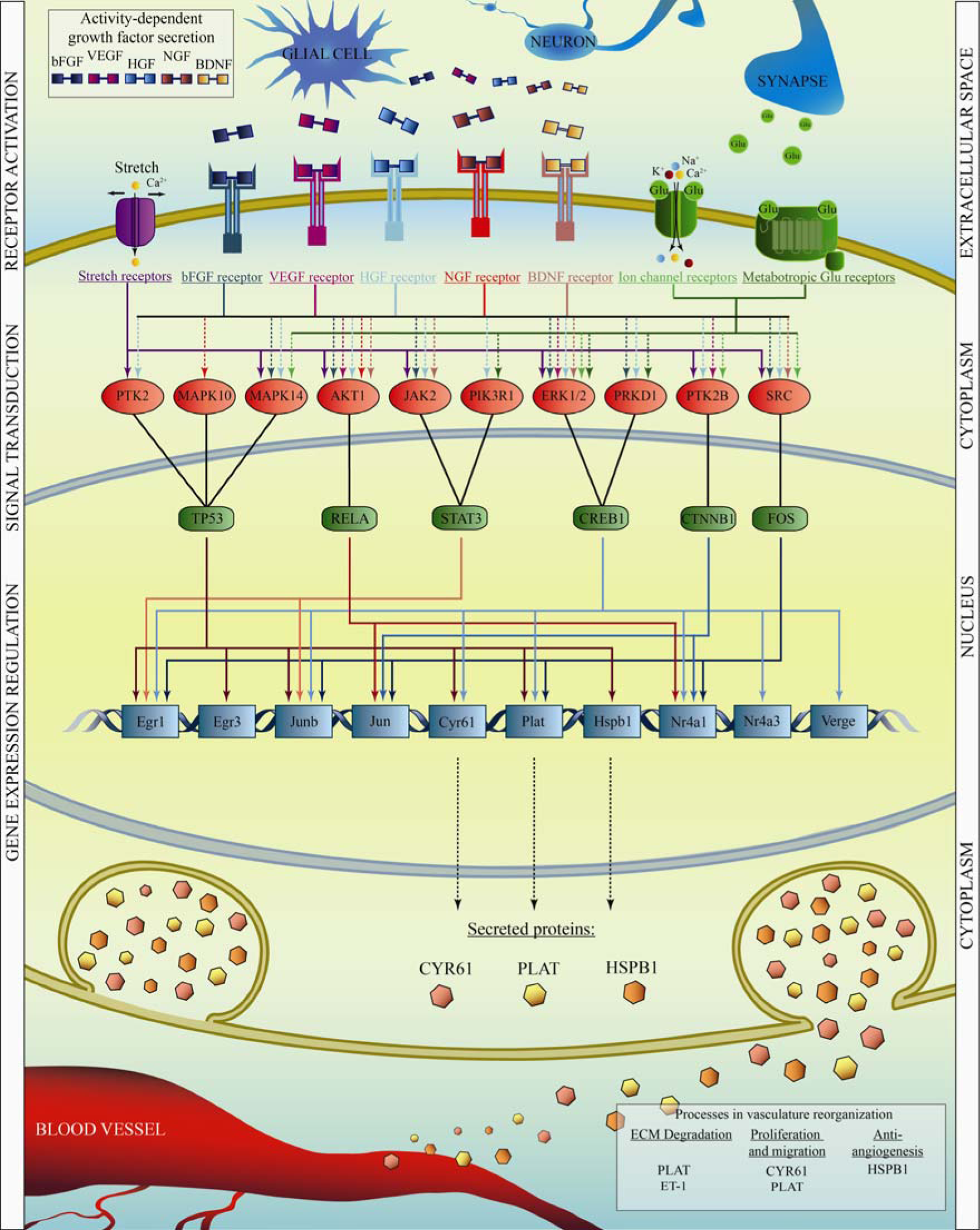
Linking synaptic activation to neurovascular reorganization. See main text for details.

A member of the heat shock proteins, *Hspb1* is ubiquitously expressed and plays important roles in apoptosis (Concannon et al., 2003) but also in smooth muscle cell migration (Hedges et al., 1999) and protection against ischemic injury (Martin et al., 1997). The latter suggests important functional roles for HSPB1 in the vasculature plausibly via its phosphorylation by VEGF (Evans et al., 2008).Modulation of the (neuro)vasculature is likely to include anti-angiogenic cues, as runaway angiogenesis is commonly associated with tumor development (De Palma et al., 2017). HSPB1 might also act as a negative regulator of angiogenesis through its interaction with VEGF upon its release from endothelial cells (Lee et al., 2012). With strong, long-lasting upregulation upon EEE, HSPB1 could orchestrate vascular responses to increased neuronal activity.

Although not most strongly regulated, several other genes among the genes upregulated upon EEE have known functions in the vasculature, see e.g molecules involved in PDGF-B signaling (**Supplemental Table 4**). PDGF-B (upregulated upon EEE; (Vallès et al., 2011)) can induce expression of *Egr1*, *Fos*, *Jun*, *Junb*, *Cyr61*, *Rap1b* and *Klf2*, (O’Brien et al., 1990; Quarck et al., 1996; Rothman et al., 1994; Wu et al., 2008) the latter of which has been shown to have, among others, *ET-1* as a downstream regulatory target (Dekker et al., 2005). ET-1, in turn, may exert its vascular functions by inducing the expression and release of vasodilatory or vasoconstrictory molecules (Hynynen and Khalil, 2006; Ihara et al., 1991). PDGF-BB-mediated induction of chemotaxis and mitogenesis in vascular smooth muscle cells requires PLAT, linking PDGF-B to yet another vasculature modulator (Herbert et al., 1997). Because these genes are expressed in an activity-dependent manner (based on (Vallès et al., 2011)), pathways such as these could well orchestrate blood vessel patterning to suit the needs of neuronal circuits (Lacoste et al., 2014; Whiteus et al., 2014).

#### 4.2) Linking synaptic activity to vasculature change through releasable molecules

The genes outlined above are shown to contribute to neurovascularization, however it is not known whether synaptic activity could regulate their transcription, and whether there are downstream protein-protein interactions that could link plasticity of synaptic communication to the organization of the neurovascular bed. Because experience dependent plasticity of neural activity is the primary mechanism by which receptive fields and sensory representations in the brain reorganize, we addressed the upstream pathways that link that these genes to the synapse. To this end, we utilized the Ingenuity^®^ Pathway Analysis (IPA^®^, Qiagen) to investigate the potential regulatory pathways in which the genes of interest are involved. In pathway reconstruction we only included experimental data from neurons, astrocytes, endothelial cells, fibroblasts, smooth muscle cells (as these three can be found in blood vessels), the CNS and CNS cell lines. Molecules were only included if they are expressed in these cell types; interactions between molecules were allowed to be indirect or direct but always filtered for activating relationships between receptors, kinases and transcription factors (e.g. kinases activating downstream transcription factors). Between transcription factors and downstream genes, relationships were filtered for ‘expression’ or ‘transcription’. Since we aimed to provide an overview of common pathways, only transcription factors that target at least two of the ten analyzed genes were used in pathway reconstruction. This approach yielded an overview of the inferred pathways that may be activated as a result of stretch, growth factor and glutamate receptor stimulation and ultimately drive expression of genes involved in remodeling of the neurovasculature (**Figure 6**).

Mechanoreceptors can be found in vasculature-related cells and astrocytes, where they can be activated as a result of increased blood flow, leading to an influx of calcium (Morita et al., 1994; Ostrow et al., 2011; Zheng et al., 2008). Among the mechanoreceptors, downstream of AGTR1 and TRPc5 (Barauna et al., 2013) (Shen et al., 2015) we find PTK2, MAPK10, MAPK14, AKT1, JAK2, PIK3R1, ERK1/2, PTK2B and SRC as downstream kinases which in turn allow transcriptional regulation of the genes of interest (**Figure 5**). Note that NMDA receptors were recently found to be activated by mechanical stress (Maneshi et al., 2017), suggesting a (partial) overlap between stretch and ionotropic glutamate receptor pathways suggested herein. In the growth factor pathway we included basic fibroblast growth factor (bFGF), hepatocyte growth factor (HGF), vascular endothelial growth factor (VEGF), brain derived neurotrophic factor (BDNF), and nerve growth factor (NGF) which show enhanced expression in an activity dependent manner in neurons and/or astrocytes (Pechán et al., 1993; Tyndall and Walikonis, 2007);(Bengoetxea et al., 2008);(Matsuda et al., 2009);(Bruno and Cuello, 2006; Pechán et al., 1993). Particularly AKT1 and ERK1/2 are downstream of these receptors. The glutamate receptor pathways include ionotrophic, α-amino-3-hydroxy-5-methyl-4-isoxazolepropionic acid (AMPA) and N-Methyl-D-aspartic acid (NMDA) receptors, activation of which lead to influx of Ca2^+^, K^+^ and/or Na^+^, as well as metabotropic glutamate receptors (mGluRs). The pathway analysis showed that glutamate receptor activation is likely to induce activation of particularly ERK1/2, PRKD1, MAPK14 and SRC kinases, leading to activation of CREB1, TP53 and FOS (**Figure 6**).

The most inclusive transcriptional regulators, suggested by this “synaptic activity to neurovascularization” pathway, are CREB1, FOS and TP53, which together can potentially regulate all our genes of interest with the exception of KLF2. *Fos* was also found by Vallès et al to be upregulated along with *Trp53rk* (TP53 regulating kinase), which increases the likelihood of FOS and TP53 involvement in the predicted pathways. Surprisingly, activating protein 1 (AP-1) was not found to have any role in any of the three pathways, even though two of its subunits were found to be upregulated after EEE (*Jun*, *Junb*) (Vallès et al., 2011) and have many potential upstream transcription factors (**Figure 6**). This suggests that the role of AP-1 either lays in the regulation of genes with vasculature-unrelated functions or that its upstream pathway involves synaptic receptors mapped herein. Lastly, no regulators were found upstream of *Prkx* and *Rap1b* that fit in the obtained pathways, suggesting that there is at least another class of upstream regulators that could link the synaptic activity to neurovascularization. Several genes (namely Egr3, Hsbp1, Nr4a3 and Verge) are downstream to only one transcription factor in our pathway. Given that the current pathways are purposely comprised of only the most prominent candidates, expression of these genes is likely regulated by other transcription factors that were excluded in the current analysis.

### 5) From synaptic plasticity to synaptic disorders

Neuronal communication deficits are likely to be central to the pathology of various brain disorders, including, but not limited to neurodevelopmental, neurodegenerative and memory disorders. Therefore, it is feasible that genes whose transcription are regulated by sensory experience might be dysregulated in select neurological disorders. In agreement with this proposition, perturbation of the plasticity-related proteins BDNF, Neuregulin 1 or mGluR1 is shown to result in a schizophrenia-like phenotype (Angelucci et al., 2005; Brody et al., 2003; O’Tuathaigh et al., 2008). Suppression of *Pak3* expression (associated with X-linked intellectual disability) in rats disrupts hippocampal plasticity (Boda et al., 2004) and when synapsin 1-3, which are involved in synaptic neurotransmitter release, are knocked out in mice, the animals display Autism Spectrum Disorder-related phenotypes of social impairments (Greco et al., 2013). While Synapsin 3 is also linked to schizophrenia-like behaviour (Porton et al., 2010), aberrant phosphorylation of eukaryotic Initiation Factor 2 (eIF2) can cause memory impairments, and its suppression reduces synaptic plasticity and memory impairments in Alzheimer’s disease model mice (Ma et al., 2013).

Moving beyond animal models, many genome-wide association (GWA) and population studies are focused on discerning the underlying genetic causes of (and predispositions to) neurological disorders. We therefore asked whether EDP-related transcripts and proteins identified in the studies by Vallès *et al* (Vallès et al., 2011) and Butko *et al* (Butko et al., 2013) are differentially regulated in neurological disorders. We used the Disease and Gene Annotations (DGA) database (Peng et al., 2013), which entails documented relationships between genes and diseases/disorders (such as altered protein or mRNA levels, single-nucleotide polymorphisms and mutations). Although the transcripts and proteins that Butko and Vallès identified as differentially regulated hardly overlap, the disorders with which they have been associated by population studies show a striking similarity, despite the divergent protein classes they encompass (**Figure 7**). This could be explained in part by the notion that these particular disorders are studied heavily.

**Figure 7.**
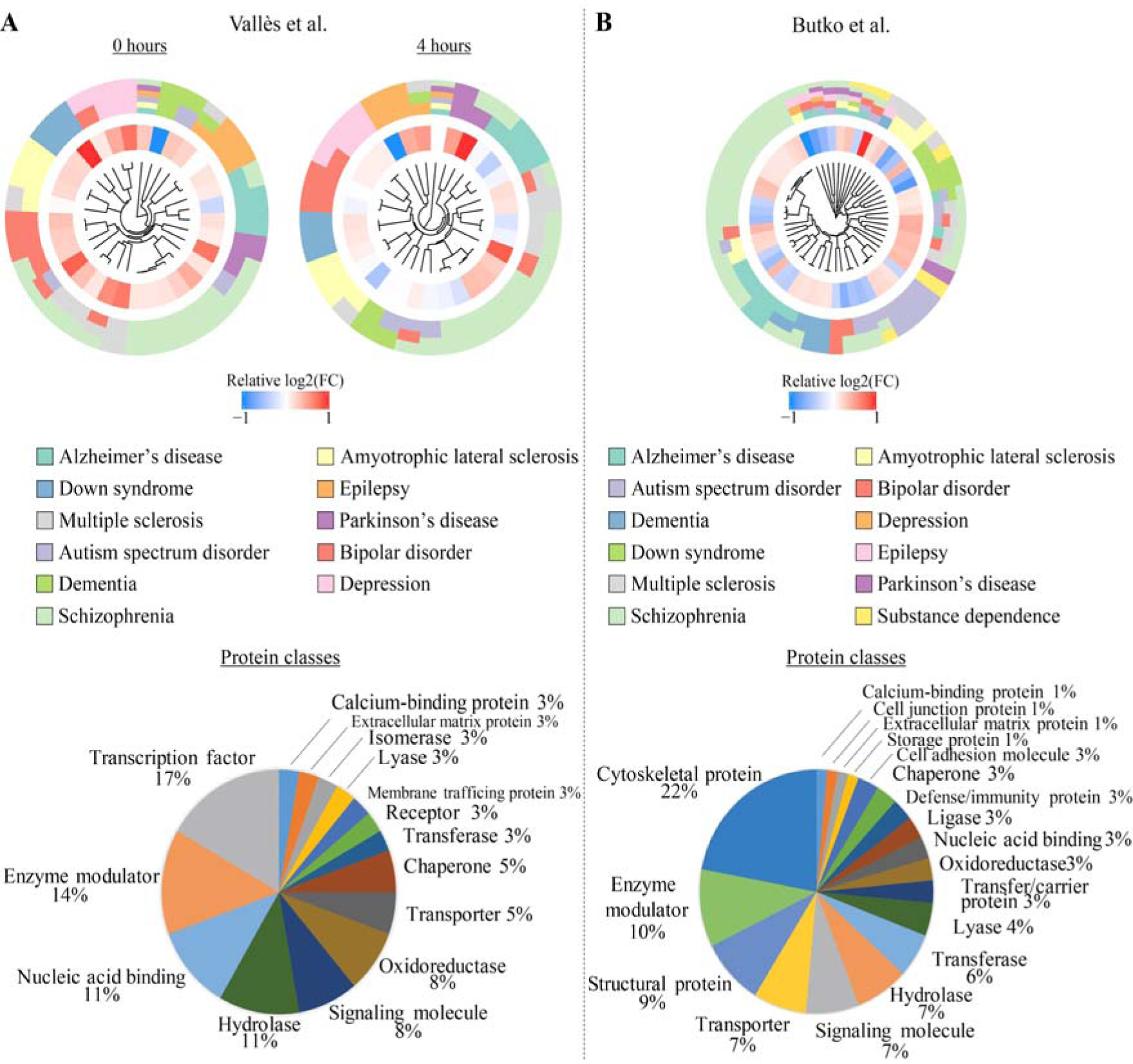
Experience-dependent plasticity and select brain disorders share common molecular targets. Top: Neurological disorders associated with differentially expressed transcripts after exposure to an enriched environment (left, data from (Vallès et al., 2011)) or proteins (right, data from (Butko et al., 2013)); genes are clustered by disorder. Bottom: Protein classes of transcripts (left) or proteins (right) associated with neurological diseases (Based on Panther Database).

Neuronal plasticity is altered in various neurological disorders, including schizophrenia (Voineskos et al., 2013), intellectual disability (Ramakers et al., 2012), autism (Pardo and Eberhart, 2007) and bipolar disorder (Schloesser et al., 2008). Targets from both molecular datasets discussed in this review can be linked to many of common neurological disorders including Alzheimer’s disease, autism spectrum disorder and schizophrenia (**Figure 7**). Some molecules are currently not known to be affected in multiple disorders, which seems to be particularly the case with transcripts or proteins related to schizophrenia, while others are affected in up to seven distinct brain disorders. For example, GRIN2B is associated with Alzheimer’s disease (Hohman et al., 2016), autism (Hu et al., 2016), bipolar disorder (Kato, 2007), Parkinson’s disease (Lee et al., 2009) and schizophrenia. Since GRIN2B is an NMDA receptor subunit, this observation likely reflects its central role in plasticity and neurological function in general. Due to the difference in sample isolation procedures in the studies by Vallès and Butko, i.e. whole tissue punch vs synaptoneurosomes, respectively, the molecules linked to brain disorders also differ substantially in the protein classes they encompass (**Figure 7**). For example, cytoskeletal proteins make up a large portion of the disease-related targets identified by Butko *et al* (Butko et al., 2013) and transcription factors are not found, whereas the reverse is true for the targets based on Vallès *et al* (Vallès et al., 2011).

An important caveat of the observations is that the links that can be established can only be as numerous as the observations that they are based on. Nevertheless, complementing GWA or population studies with experimental evidence such as those obtained from transcriptomics or proteomics studies can facilitate identification of underlying causes of neurological disorders or understand how symptoms during disease progression arise. Although transcriptome- or proteome-wide studies provide direction for research, in particular when combined with expression data obtained from patients, functional studies remain indispensable to fully understand the molecular bases of health and disease. As experience dependent plasticity and a large number of brain disorders involve similar transcriptional and translational targets, experience dependent plasticity in sensory circuits could help to unravel the molecular pathways associated with brain disorders.

### 6) Future directions for molecular studies of experience-dependent plasticity

Transcriptomic and proteomic big data have an outstanding promise to usher the field of experience-dependent plasticity to a new era where differential transcriptional regulation of genes are linked to neural network reorganization through large-scale molecular pathway analysis, causally linking molecules to network organization and ultimately to behaviour.

EDP requires the interaction of the numerous different cell types that exist in the brain, from excitatory (Allen et al., 2003) and inhibitory (Foeller et al., 2005) neurons to glia (Perez-Alvarez et al., 2014) and even the epithelial and muscle cells of blood vessels (Lacoste et al., 2014; Whiteus et al., 2014). A major unanswered question is how the vastly diverse cortical cellular population orchestrates EDP and which of the molecules that underlie it are common across (all) cell types and which are critical to a select few. Current technologies allow for the isolation of individual cells, possibly in combination with fluorescent labeling of molecularly defined cell classes, from which RNA (Zeisel et al., 2015) or (although still in need of development) proteins (Heath et al., 2016) can be harvested and subsequently analyzed. Using currently available datasets, it is possible to infer *post hoc* which cell types most robustly changed their transcriptomic make-up following sensory experience, as we have exemplified above. Nonetheless, applying single-cell technologies in combination with sensory experience manipulations would enable the study of molecules critical for EDP with unprecedented precision.

A further benefit would come from the collection of samples in a cortical layer-specific manner. Electrophysiologically, cortical lamina have since long been shown to display distinct phenotypes in cortical map plasticity (Diamond et al., 1994), stemming from the organization of their thalamic and cortical inputs and their cellular populations (Molyneaux et al., 2007). Cortical laminae can be readily identified through molecular markers, showing their distinct molecular identities (Belgard et al., 2011; Molyneaux et al., 2007), which likely shows the coherence between a layer’s functional role and the molecules required to exert it. This stresses the importance of anatomical resolution when studying experience-dependent plasticity on either the electrophysiological and molecular level. Both Vallès (Vallès et al., 2011) and Butko (Butko et al., 2013) and their colleagues have examined, through *in situ* hybridization or immunostainings, the distribution of a subset of identified targets across cortical lamina, revealing tight spatial restrictions on their expression in cellular populations. Such *post-hoc* approaches however are of low throughput, and future studies should strive to obtain lamina-specific tissues (much like in Belgard et al., 2011; Kole et al., 2017a, 2017b) and use these for subsequent processing and analysis. This will allow for a large-scale screening of layer-specific manifestations of experience-dependent plasticity at the molecular level.

The most recent transcriptome and proteome studies employ environmental enrichment or all-whisker stimulation or trimming, which induce homeostatic plasticity in barrel cortex. Homeostatic and Hebbian plasticity work in coherence to establish cortical maps, but their mechanisms differ significantly (Feldman and Brecht, 2005; Gainey and Feldman, 2017; Keck et al., 2017). Alternative deprivation or stimulation methods, such as single-row whisker deprivation or single-whisker experience (Allen et al., 2003; Celikel et al., 2004; Clem et al., 2008; Foeller et al., 2005), result in electrophysiologically well-defined plasticity phenotypes, the molecular mechanisms of which have been poorly studied and still require elucidation.

The ultimate goal will have to be to systematically study experience-dependent plasticity in a cell, cell-type, node (e.g. cortical layer) specific manner, across the many forms of experience (see **Figure 2**) and perceptual learning-induced. This will allow researchers to create an all-inclusive molecular map of the transcriptome and proteome, that will lead to causal experiments to unravel the molecules that control brain plasticity plasticity (Kole et al., 2017a; Kole et al., 2017b). The whisker system, with its modular organization along the whisker-to-barrel pathway as well as the rich behavioral repertoire of whisker dependent tasks (e.g. (Filipkowski et al., 2000; Guic-Robles et al., 1992; Jabłońska and Skangiel-Kramska, 1995; Jacobs and Juliano, 1995);(Celikel and Sakmann, 2007; Clem et al., 2008; Cohen and Castro-Alamancos, 2010; Galvez et al., 2007; Grion et al., 2016; Guic et al., 2008; Juczewski et al., 2016; Krupa et al., 2001; Miceli et al., 2017; Miyashita and Feldman, 2013; O’Connor et al., 2010; Rosselet et al., 2011; Safaai et al., 2013; Topchiy et al., 2009; Troncoso et al., 2007; Voigts et al., 2015; Waiblinger et al., 2015)), will likely lead the way.

## Acknowledgements

This work was funded by the Faculty of Science of the Radboud University, Nijmegen, the Netherlands (grant number 626830 – 6200821) and the ALW Open Programme of the Netherlands Organization for Scientific Research (NWO; grant number 824.14.022).

## Supplemental material

### Supplemental Table 1

Differentially expressed genes of GO terms shown in Figure 3 (data is from Vallès et al., 2011).

### Supplemental Table 2

Functional clusters of Gene Ontology (GO) terms for the differentially expressed genes in the Vallès et al. (2011) dataset. Only clusters with an FDR of <0.05 and an enrichment score of >1.3 are included. The most significant term (with the lowest FDR) within each cluster is displayed.

### Supplemental Table 3

Cellular Enrichment Indices (CEIs) of differentially expressed genes upon EEE. See Figure 4 for the details of the bioinformatics pathway.

### Supplemental Table 4

List of differentially expressed transcripts within the GO term ‘Vasculature Development’ (see Supplemental Table 2). In bold are the transcripts that are most strongly regulated upon EEE and remain differentially transcribed for at least 4 hours (cutoff 10%). Transcripts in italics are those involved in PDGF-B signaling.

